# Direct detection of circulating microRNA-122 using dynamic chemical labelling with single molecule detection overcomes stability and isomiR challenges for biomarker qualification

**DOI:** 10.1101/777458

**Authors:** Barbara López-Longarela, Emma E. Morrison, John D. Tranter, Lianne Chahman-Vos, Jean-François Léonard, Jean-Charles Gautier, Sébastien Laurent, Aude Lartigau, Eric Boitier, Lucile Sautier, Pedro Carmona-Saez, Jordi Martorell-Marugan, Richard J. Mellanby, Salvatore Pernagallo, Hugh Ilyine, David M. Rissin, David C. Duffy, James W. Dear, Juan J. Díaz-Mochón

## Abstract

Circulating microRNAs are biomarkers reported to be stable and translational across species. miR-122 (miR-122-5p) is a hepatocyte-specific microRNA biomarker for drug-induced liver injury (DILI). Our objective was to develop an extraction-free and amplification-free detection method for measuring miR-122 that has translational utility in context of DILI. We developed a single molecule dynamic chemical labelling (DCL) assay based on miR-122 hybridization to an abasic peptide nucleic acid probe that contained a reactive amine instead of a nucleotide at a specific position in the sequence. The single molecule DCL assay specifically measured miR-122 directly from 10 µL of serum or plasma without any extraction steps, with a fit-for-purpose limit of detection of 1.32 pM. In 192 human serum samples, DCL accurately identified patients at risk of DILI (area under ROC curve 0.98 (95%CI 0.96-1), P<0.0001). The miR-122 assay also quantified liver injury in rats and dogs. When DCL beads were added to serum, the miR-122 signal was stabilised (no loss of signal after 14 days at room temperature). By contrast, there was substantial degradation of miR-122 in the absence of beads (≈60% lost in 1 day). RNA sequencing demonstrated the presence of multiple miR-122 isomiRs with DILI that were at low concentration or not present in healthy patient serum. Sample degradation over time produced more isomiRs, particularly rapidly with DILI. PCR was inaccurate when analysing miR-122 isomiRs, whereas the DCL assay demonstrated accurate quantification. In summary, the DCL assay can accurately measure miR-122 directly from serum and plasma to diagnose liver injury in humans and other species, and can overcome important microRNA biomarker analytical and biological challenges.

## Introduction

MicroRNAs are non-protein coding RNAs consisting of approximately 22 nucleotides. They have a key role in post-transcriptional regulation of gene expression in diverse biological processes such as cell differentiation and proliferation, metabolism and apoptosis.^1^ Dysregulation of microRNA expression is associated with a wide range of diseases such as cancer, infections and metabolic disorders. Cell-free microRNAs are present in biological fluids where their concentration can change when disease states develop.^2^ In blood, microRNAs are reported to be relatively stable because they are protected from enzymatic degradation by being encapsulated in extra-cellular vesicles (ECVs) or bound to protein complexes.^3^ These properties make microRNA a promising new class of disease biomarker, which is translational due to microRNA sequences being highly conserved across species.

However, there are roadblocks that prevent microRNA species being adopted as biomarkers in clinical practice. The gold standard method for quantification of circulating microRNA is RT-qPCR. Technical challenges for using PCR in this context include pre-analysis degradation of the target microRNA, variability in the extraction of microRNAs from plasma or serum, matrix effects exemplified by heparin inhibition of polymerases, operator variability and a lack of consistency across studies regarding the optimal methods used to normalize data. Furthermore, PCR is relatively expensive both in terms of financial cost and need for technically skilled personnel, and may be too slow to provide the rapid turnaround results needed to inform acute clinical decision-making.

Contrary to initial belief, during microRNA biogenesis a microRNA precursor does not produce a single mature microRNA. Instead, precursors produce a ‘cloud of isoforms’ known as isomiRs.^4, 5^ isomiRs from the same precursor microRNA differ slightly in their lengths at the 5’ termini, 3’ termini, or both. These isomiRs increase the complexity of correctly identifying and quantifying specific microRNA species. Specific isomiR patterns have been reported to stratify diseases, such as cancer^6^, suggesting that accurate measurement of specific isomiRs may have clinical utility. At present, RNA sequencing is the only technique able to accurately detect and quantify isomiRs.

The adoption of microRNA into clinical practice would be galvanized by the development of a simple, direct, quantitative, and reproducible testing protocol for the detection of circulating microRNAs with single base resolution, a fit-for-purpose sample-to-result turnaround time, and relative operator independence. To address this challenge, we have developed a PCR-free, sensitive and specific assay for measuring microRNA.^7^ This dynamic chemical labelling (DCL) assay is based on microRNA hybridization to an abasic peptide nucleic acid probe—containing a reactive amine instead of a nucleotide at a specific position in the sequence—attached to superparamagnetic beads. This step is followed by the specific incorporation of a biotinylated reactive nucleobase. Our previous publication demonstrated that this approach can sensitively and specifically quantify microRNA targets isolated from human serum.^7^ In this paper we determine its utility with regard to direct detection in a range of pre-clinical and clinical samples.

MicroRNA-122 (miR-122, miR-122-5p) is a microRNA species that is highly expressed in, and highly specific for, hepatocytes, with little to no expression in other tissues (∽40,000 copies per hepatocyte).^8^ In liver injury, miR-122 is released from necrotic hepatocytes, resulting in substantially elevated miR-122 concentrations in the bloodstream.^2^ In patients with acute liver injury, we reported that the circulating concentration of miR-122 is increased 100-1000 fold compared to controls.^9^ In over 1000 patients, we recently demonstrated in the Markers and Paracetamol Poisoning (MAPP) Study that miR-122 is a sensitive and specific biomarker of acute liver injury after paracetamol overdose (the commonest cause of drug-induced liver injury (DILI) in the Western world).^10^ miR-122 also has potential to aid the process of preclinical and clinical drug development. This point was acknowledged by regulatory support from the US Food and Drug Administration (FDA) and the European Medicines Agency (EMA) for miR-122 to be used as an exploratory DILI biomarker in clinical trials. However, the clinical development of miR-122 as a biomarker has been challenged because the reference interval in healthy subjects is wide when miR-122 is measured using standard PCR techniques.^11^ It is vital for the field to understand whether this variability is due to biological or analytical factors.

In this study we have used circulating miR-122 and DILI as the clinical paradigm to test whether the DCL assay can sensitively and specifically measure microRNA directly from serum or plasma with no need for microRNA extraction or amplification. The application of this new analytical approach to microRNA measurement facilitated the exploration of key translational questions regarding microRNA stability, degradation, and the effect of isomiRs on microRNA quantification.

## Material and Methods

### Materials

RNA target molecules were purchased desalted from Integrated DNA Technologies. 2.8-µm diameter superparamagnetic beads presenting carboxylic acid groups (Dynabeads® M-270), were obtained from ThermoFisher Scientific. Streptavidin-β-galactosidase (SβG), resorufin-β-D-galactopyranoside (RGP), Simoa disks, the Simoa microplate shaker, the Simoa Washer and SR-X reader were obtained from Quanterix Corporation. Abasic PNA probes, superparamagnetic capture probe beads (DGL-122_4.3) and biotinylated reactive cytosine nucleobase were prepared as reported elsewhere.^7^ DestiNA Stabiltech Buffer is a lysis buffer containing lithium salts and detergent. DestiNA Reaction Buffer is made of 10% w/v PEG-10K, in a buffered solution of 30 mM trisodium citrate and 300 mM sodium chloride, pH adjusted to 6.0 using HCl. Concentrations of solutions of RNA and DNA were determined using a ThermoFisher NanoDrop1000 Spectrophotometer.

### Collection and preparation of clinical samples

Patient serum samples collected in the MAPP study were analysed. The full details of this study are published elsewhere. ^10^ In brief, adult patients (16 years old and over) were recruited if they fulfilled the study inclusion and exclusion criteria. Full informed consent was obtained from every participant and ethical approval was given by the South East Scotland Research Ethics Committee and the East of Scotland Research Ethics Committee via the South East Scotland Human Bioresource. The inclusion criteria were: a history of paracetamol overdose that the treating clinician judged to warrant treatment with the antidote (intravenous acetylcysteine) as per the contemporaneous UK guidelines; the first blood sample collected within 24 hours of last paracetamol ingestion and the patient had capacity to consent. The exclusion criteria were detention under the Mental Health Act; documented cognitive impairment; inability to provide informed consent for any reason or an unreliable overdose history. Demographic information was recorded for the study participants and their blood sample taken at first presentation to hospital was stored at −80°C as plasma/serum. The primary endpoint for the MAPP Study was acute liver injury, pre-defined as a peak hospital stay serum alanine transaminase activity (ALT) greater than 100U/L.

### Collection and preparation of animal samples

#### Rat model of drug-induced liver injury

An investigative rat toxicology study was conducted. In brief, purpose-bred male Sprague Dawley rats (age at initiation of dosing: 9-10 weeks) were supplied by Charles River Laboratories (Calco, Italy) and acclimatized for 14 days before the start of the experiment. The animals were housed in a facility accredited by the Association for Assessment and Accreditation of Laboratory Animal Care International. The study and all procedures were in accordance with the Directive 2010/63/EU of the European parliament and the related French transposition texts, and were approved by the Sanofi Animal Care and Use Committee. The day of arrival, the animals were randomly allocated to the vehicle group (6 rats) or the compound-dosed group (6 rats). The 6 animals from the compound-dosed group were given a single oral administration of paracetamol in the vehicle (Methylcellulose, Tween 80, Water at 0.5, 0.1 and 99.4%, respectively) at the dose level of 1500 mg/kg, and the control group received the respective vehicle. Food was withheld overnight before necropsy. 24 hours after single dosing, animals were anesthetised with isoflurane and blood samples were collected at the abdominal aorta for clinical chemistry analyses (900 μL in heparin lithium vacutainer tubes) and miR-122 measurements using the DCL and droplet-digital PCR (ddPCR) assays (400 μL in EDTA K2 vacutainer tubes frozen and stored at −80°C). Animals were then euthanized by exsanguination and a full necropsy was performed with macroscopic examination of the thoracic and abdominal cavities and organs. The liver, heart, kidneys and brain were weighed. The liver (left and right lateral lobes as well as median lobe), heart, kidneys and *quadriceps femoris* muscle were collected for histopathological evaluation, fixed in 10% neutral buffered formalin, and paraffin-embedded within 48 hours. Sections were then stained with haematoxylin and eosin (H&E) and examined by two board-certified veterinarian pathologists. Microscopic findings were classified according to the INHAND (International Harmonization of Nomenclature and Diagnostic Criteria) nomenclature system ^12^ and graded on a scale of 0–5, where 0 = no change, 1 = minimal, 2 = slight, 3 = moderate, 4 = marked and 5 = severe. Plasma alanine transaminase (ALT) and glutamate dehydrogenase (GLDH) activites were measured on a Roche Cobas 6000 c501 on an automated clinical chemistry analyser using reagents purchased from Roche Diagnostics.

#### Dogs with clinical liver disease

All dogs were recruited to this study at the Royal [Dick] School of Veterinary Studies [R[D]SVS], Edinburgh, UK. Healthy dogs presenting to the R[D]SVS General Practice for routine annual vaccination, who had a normal history and clinical examination, were invited to have a serum biochemical health screen which included measurement of ALT activity. Dogs that had a diagnostic assessment that included histopathological examination of a liver biopsy and serum ALT activity measurement leading to definitive diagnosis of a primary liver disorder were also enrolled into the study. The histopathological diagnosis was classified according to WSAVA [World Small Animal Veterinary Association] criteria by a board certified pathologist. The study was approved by The University of Edinburgh Veterinary Ethics Research Committee.

#### DCL Assay

The DCL assay was performed in two steps: (i) Hybridization – serum samples were used immediately after collection, with addition of Stabiltech Buffer and capture probe beads DGL-122_4.3, or frozen and stored at −80°C for later use (probe sequences in **Supplementary Table 6**). Frozen samples were first thawed at room temperature. Serum samples (10 µL) were added to 25 µL of Stabiltech Buffer containing 100,000 DGL 122_4.3 beads placed in a 96 well plate and were mixed by pipetting. Annealing was performed by rotating in a Simoa microplate shaker at 800 rpm for 1 h at 25°C. Samples were analysed immediately, or in specified experiments, kept at room temperature for up to 28 days. For the calibration curve, different concentrations of ssmiR-122 in the same matrix as biological samples were prepared. Following 1h hybridization step, the DGL 122_4.3 beads were washed three times with post-hybridization washing buffer. (ii) Dynamic Chemistry Labelling (see **Supplementary Figure 5** for the chemical reaction) – After the washing steps, 50 µL of master mix was added to each well. Master mix was a solution of 5 µM aldehyde-modified biotinylated cytosine and 1 mM sodium cyanoborohydride in reaction buffer. The microtiter plate was placed on an incubating shaker, set to 40°C, and mixed at 800 rpm for 1 h. Then beads were washed three times with washing buffer followed by resuspension of the beads in 100 μL of 25 pM SbG and incubation for 10 min at 30°C on the microplate shaker set at 800 rpm. Following the 10 min incubation time, beads were washed and analysed on the SR-X Reader to determine the active enzyme per bead (AEB) values. The SR-X enables reading of single enzyme labels on the beads using the Simoa technology.^13^ The assay steps and single molecule detection were performed separate from Simoa detection, allowing the DCL assay steps to be performed on standalone washer and incubators using customized buffer compositions.

#### Extraction of microRNA from serum samples

microRNA was extracted using a miRNeasy Serum/Plasma kit [Qiagen, Venlo, Netherlands] following the manufacturer’s instructions. Total RNA was extracted from 50 μL of serum diluted in 150 μL nuclease free water. Briefly, RNA was extracted from the serum by lysis reagent [1000 μL] and chloroform [200 μL]. After centrifugation at 12,000 x g for 15 min at 4°C up to 600 μL of the aqueous phase was transferred to a new tube with 900 μL absolute ethanol. RNA was purified on an RNeasy minElute spin column and eluted in 15 μL RNase-free water and stored at −80°C. Extraction efficiency was monitored by adding 5.6 × 10^8^ copies of synthetic *C.elegans* miR-39 spike-in control after the addition of lysis reagent before the addition of chloroform and phase separation.

#### Reverse Transcription and Real-Time Polymerase Chain Reaction [RT-PCR]

##### SYBR Green PCR

The miScript II Reverse Transcription Kit [Qiagen, Venlo, Netherlands] was used to prepare cDNA according to the manufacturer’s instructions. Briefly, 2.5 μL of RNA eluate was reverse transcribed into cDNA. The synthesised cDNA was then diluted and used in combination with the miScript SYBR Green PCR kit [Qiagen] and a specific miScript assay targeting hsa-miR-122-5p (Hs_miR-122a_1) [Qiagen] for qPCR. qPCR was performed on the LightCycler 480 [Roche, Burgess Hill, UK] system using cycling parameters recommended for miScript assays. All samples were ran in triplicate.

##### Taqman PCR

RNA eluates were reverse transcribed by following the manufacturer’s instructions with the TaqMan MicroRNA Reverse Transcription Kit [Applied Biosystems, Foster City, CA, USA] and TaqMan MicroRNA assay hsa-miR-122 (Assay ID: 002245) [Applied Biosystems]. In reverse transcription, 5 μL of RNA was transcribed to complementary DNA (cDNA) in a total volume of 15 μL. Then, 1.33 μL of cDNA was used in combination with the TaqMan Universal PCR Master Mix, no AmpErase UNG [Applied Biosystems] and TaqMan probe from the hsa-miR-122 assay [Applied Biosystems] in a total volume of 20 μL. qPCR was performed on the LightCycler 480 [Roche] system using cycling parameters recommended for TaqMan assays. All samples were ran in triplicate.

For SYBR Green and Taqman PCR Serial dilutions of known standards were made using synthetic miR-122 (syn has-miR-122-5p, 219600, Qiagen). The dilutions were prepared in triplicate, using the same RT and qPCR protocols as described above. The Ct values were plotted against the logarithm of the concentration, demonstrating a clear linear relationship between Ct value and Log (conc.). The resultant regression line was used to ascertain the concentration of the unknown sample set.

##### Droplet-digital PCR (ddPCR)

ddPCR was performed on plasma of rats. The extraction of microRNAs was performed with a Qiacube instrument (Qiagen) using 200 μL of plasma. The miRNeasy mini kit (QIAGEN, USA) was used with phase lock gel (QuantaBio, USA) and following the last silica gel column elution, 40 μL of extracts were collected and stored at −80°C until downstream analysis. The same extraction procedure was used for spiking experiments with a synthetic rno-miR-122-5p in pooled plasma of naïve rats to evaluate extraction yield. After thawing, the reverse transcription of 5 μL of extracts was carried out using the TaqMan™ MicroRNA Reverse Transcription Kit (ThermoFisher, USA) including the TaqMan RT 5 x primer (assay ID: 002245) specific to rno-miR-122-5p; then, the cDNA were stored at −80°C. The day of ddPCR experiment, 2 dilutions (1/10 and 1/100) of cDNA were performed and the droplets were generated using the ddPCR™ Supermix, the miR122-5p specific TaqMan assay, the droplet generation oil for Probes and the QX200™ Droplet Generator (BIORAD, USA). The droplets were collected in 96 sealed well plates and run for PCR. A 40 amplification cycle at 56°C was performed with the T100™ Thermal Cycler (BIORAD, USA) then, the samples were kept at +4°C until processed the same day by a QX200 Reader (Biorad, USA). The final quantification of miR-122-5p copies was done using the QuantaSoft™ Analysis Pro software (BIORAD, USA) according to the Poisson law; all samples were evaluated in triplicate.

#### Stability study protocol

Testing was performed in two laboratories, at the University of Edinburgh Medical School, and DestiNA Genomica, Granada Spain. All the experiments were conducted on the same dates at the two locations with an identical blood serum sample prepared from a single blood sample taken from a patient presenting at the Royal Infirmary of Edinburgh Emergency Department in Edinburgh, with DILI due to paracetamol overdose (ALT>1000U/L). See Supplementary Information for full details of the study.

#### RNA sequencing

Total RNA was extracted as mentioned above. Standard quality control steps were included to determine total RNA quantity and quality using Agilent 2100 Bioanalyzer with Eukaryote Total RNA Pico Kit and High Sensitivity DNA Assay (Agilent Technologies, Santa Clara, CA). Libraries were prepared using the TruSeq Stranded mRNA Library preparation Kit (Illumina Inc., San Diego, USA) and RNA sequencing was performed on an Illumina NextSeq 500 sequencing platform.

#### Data Processing and Statistical Analysis

RNA-Seq data analysis was performed using QuickMIRSeq suite.^14^ Briefly, raw sequences were trimmed in order to remove adapter sequences and random nucleotides introduced during the library preparation. Reads shorter than 15 nucleotides (nt) and larger than 28 nt after trimming were discarded to select those sequences more likely to map against miRNAs. Reads were aligned to human reference genome GRCh38. We configured QuickMIRSeq to map not only against reference miRNAs, but also against possible isomiRs with length variations of ± 4 nucleotides at the 5’ end and ± 5 nucleotides at the 3’ end. We normalized the miRNA counts by reads per million (RPM) method in order to correct for differences in sequencing depth. We discarded low-expressed miRNAs (less than 50 RPM in at least 6 samples). We used pheatmap R package for generating heatmaps.

Data are presented as median (IQR) unless specifically indicated. Statistical analysis was as described in the results.

## Results

The experimental protocol for the DCL assay is presented in **Figure 1A.** Beads coated with miR-122 specific probes (see **Supplementary Table 6** for details of their sequence) were added to healthy human serum, with synthetic miR-122 or miR-21 spiked in, then binding to the miR-122 complementary probe was quantified without any RNA extraction or amplification. Synthetic miR-122 (see Supplementary Information for details in the sequence) could be quantified with the characteristics described in **Table 1**.

**Table 1.**
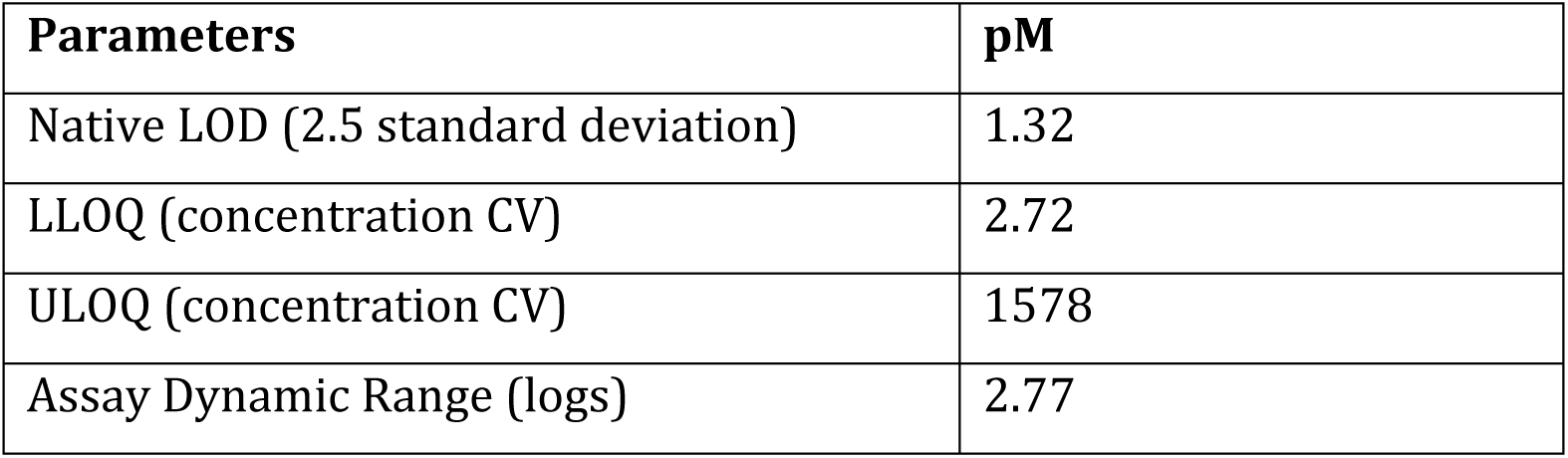
Performance characteristics of the DCL assay. Data calculated from four calibration curves. LOD = limit of detection. LLOQ = lower limit of quantification. ULOQ = upper limit of quantification.

| Parameters | pM |
| --- | --- |
| Native LOD (2.5 standard deviation) | 1.32 |
| LLOQ (concentration CV) | 2.72 |
| ULOQ (concentration CV) | 1578 |
| Assay Dynamic Range (logs) | 2.77 |

**Figure 1.**
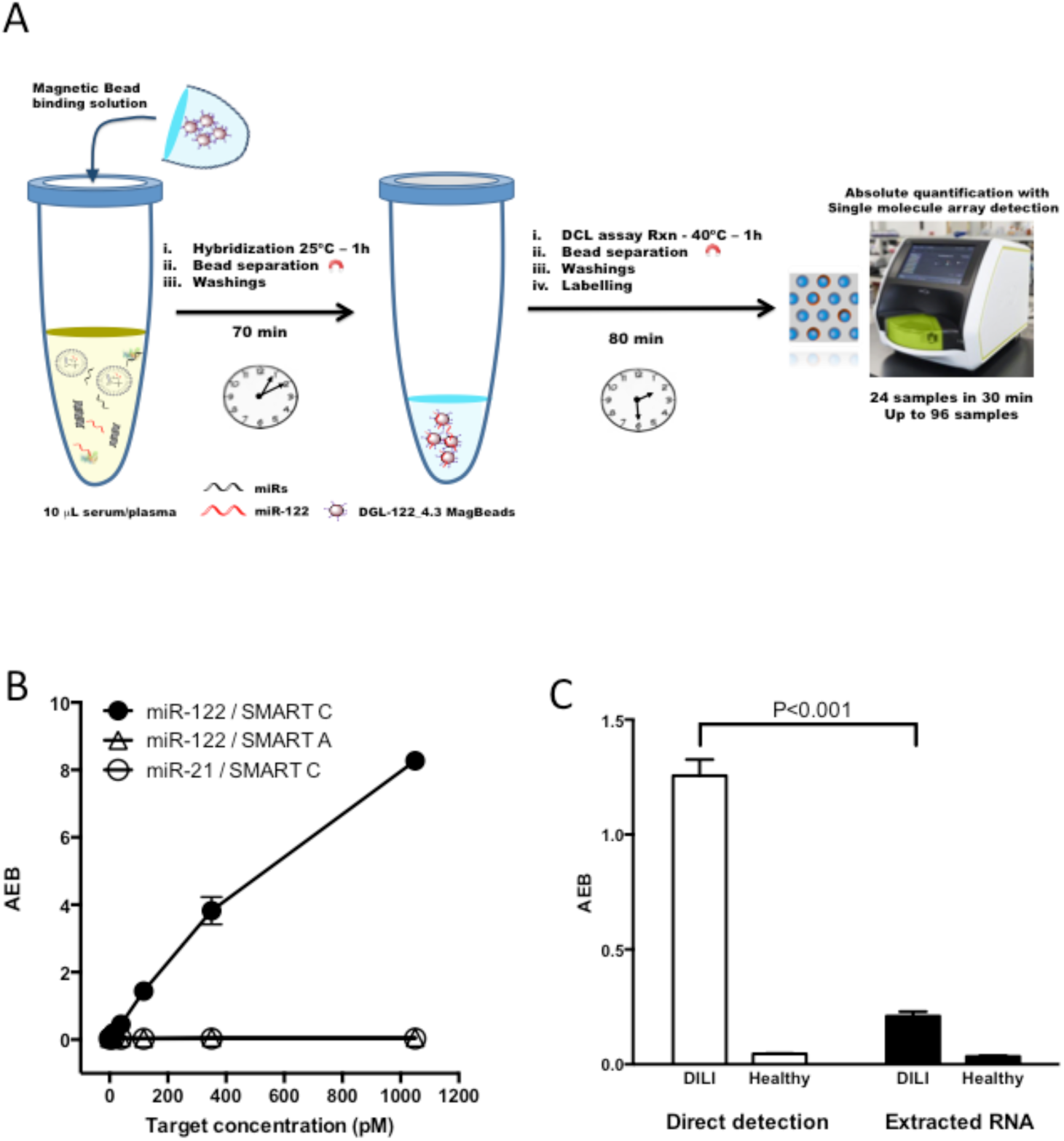
Direct detection and quantification of microRNA in serum by dynamic chemistry labelling (DCL assay) using single molecule arrays. A) DCL protocol. B) Plot of Active Enzyme per Bead (AEB) against the concentration of miR-122 or miR-21 spiked into healthy human serum. The biotinylated reactive nucleobase added was cytosine (C) or adenine (A). Error bars (±1 s.d.) are generated from triplicate measurements. C) miR-122 quantification by DCL assay using the extraction-free protocol described in panel A (Direct detection) or RNA extracted from serum as per PCR protocol (Extracted RNA). Error bars (±1 s.d.) based on triplicate measurements. P value calculated by T-Test.

There was no signal with miR-21, or with miR-122 when the biotinylated reactive nucleobase was adenine rather than the correct base (cytosine) **(Figure 1B).** Direct detection of miR-122 from human serum (with no RNA extraction step) was compared with miR-122 quantified following microRNA extraction. Serum from a patient with DILI was compared to a healthy volunteer. The fold change (DILI to health) in signal (AEB) was 27 when miR-122 was detected directly in serum compared to 6 fold following RNA extraction **(Figure 1C).**

To determine the performance of the DCL assay as a potential clinical diagnostic assay we analysed samples from the MAPP study. The MAPP study prospectively recruited patients at risk of DILI due to taking a paracetamol overdose that required treatment with the antidote acetylcysteine. In derivation and validation cohorts, this study reported that miR-122, measured by Taqman-base PCR, predicted acute liver injury (pre-defined as peak ALT activity >100 U/L) at hospital presentation, with high sensitivity and specificity (area under ROC curve 0.97 (95%CI: 0.95-0.98), sensitivity 0.79 (95%CI: 0.70-0.87) at 0.95 specificity).^10^ In the current study, we used 192 sequential serum samples from 2 hospital sites in the MAPP study to determine whether miR-122 retained high sensitivity and specificity when measured using the DCL assay. Demographic and laboratory data are presented in **Supplementary Table 1.** There was a significant correlation between the concentration of miR-122 measured by DCL and PCR **(Supplementary Figure 1).** Patients who subsequently developed acute liver injury were separated from those patients without injury by DCL **(Figure 2).** Using the DCL assay, the concentration of miR-122 was higher in those study participants who developed liver injury, compared to those who did not (no injury median 1.4 pM (IQR 0.6 - 2.7); acute liver injury 35.8 pM (15.3 - 128)). ROC analysis demonstrated that sensitivity and specificity was comparable to using PCR (area under ROC curve 0.98 (95%CI: 0.96-1), sensitivity 0.83 (95%CI: 0.59-0.96) at 0.95 specificity).

**Figure 2.**
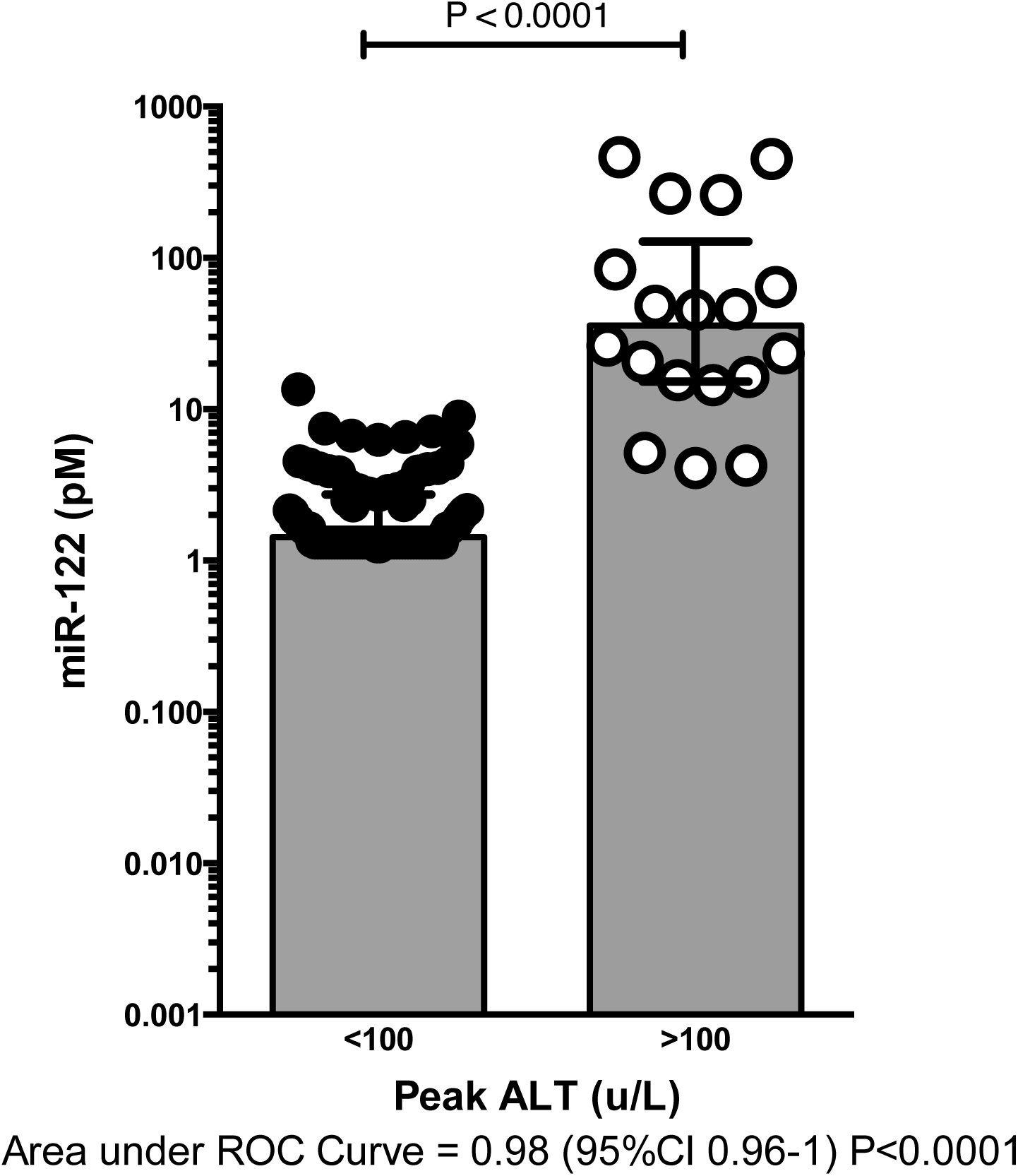
Measurement of miR-122 using the DCL assay can accurately identify patients who will subsequently develop acute liver injury at first presentation to hospital after paracetamol overdose. In 192 participants from the MAPP study, the DCL assay was used to measure serum miR-122. Acute liver injury was pre-defined in MAPP as a peak serum alanine transaminase (ALT) activity of greater than 100 U/L. Each point represents a participant, the bars represent the median, and error bars the inter-quartile range. Values below the LOD are reported as the LOD value (1.32 pM). The groups were compared by Mann-Whitney test and using a receiver operator characteristic (ROC) curve.

MicroRNAs are conserved across species (translational biomarkers) and miR-122 accurately reports liver injury in a range of models.^15^ In a rat model of paracetamol-induced liver injury, treated animals had increased plasma ALT activity 24 hours post-dose (mean = 518.8 U/L, SD = 754.2) when compared to controls (mean = 35.8 U/L, SD = 27.4 p < 0.05, Mann-Whitney test), and increased plasma GLDH activity (treated: mean = 207.7 U/L, SD = 176.0; controls: mean = 3.7 U/L, SD = 1.9, p < 0.05, Mann-Whitney test). At terminal sacrifice, the mean absolute liver weight from the treated group was 16% higher (p < 0.05, Mann-Whitney test) when compared with control animals. Microscopically, the liver from paracetamol-treated rats showed minimal to marked hepatocellular centrilobular apoptosis and/or necrosis, together with minimal to moderate multifocal neutrophilic and/or mononuclear cell inflammatory infiltration. There were no compound-related microscopic changes in the heart, kidneys and skeletal muscle from treated rats. miR-122 concentration was increased when measured in 10 µL of plasma from rats treated with paracetamol using the DCL assay (vehicle: median 8.4 pM (IQR 6 – 15.3); paracetamol: 825.1 pM (335-3365)). There was overall a correlation between miR-122 concentrations and both plasma ALT and GLDH activities and the severity of histopathological lesions in the liver **(Supplementary Table 2).** ddPCR was used to compare the results obtained with the DCL assay. There was a strong linear relationship **(Supplementary Figure 2)** between the DCL assay and ddPCR. However, when copy numbers of miR-122 per μL for each sample were compared, the ddPCR values were around three orders of magnitude lower than those calculated by the DCL assay **(Supplementary Table 2).** To further explore this discrepancy, three concentrations of miR-122 calibrator at 10 pM, 100 pM and 500 pM in naïve male rat plasma were measured by ddPCR and DCL (**Supplementary Table 3).** Values obtained by ddPCR were again more than three orders of magnitude lower than expected, while concentrations calculated by the DCL assay were as expected.

We compared serum from healthy dogs to dogs with a range of biopsy diagnosed liver diseases (n = 12 per group). The clinical characteristics of the liver disease group are presented in **Supplementary Table 4** Consistent with our previous published data using PCR ^16^, miR-122 was significantly higher in the liver disease group when measured using the DCL assay **(Supplementary Figure 3).**

To explore the effect of adding DCL assay beads on the stability of miR-122, we used serum from a patient with DILI (ALT > 1000 U/L after paracetamol overdose), which was stored at room temperature, with serial measurements of miR-122 by both DCL assay and PCR **(Figure 3).** Blood was drawn, then immediately processed to separate the serum, and frozen at −80°C. On day 0 of the experiment, the serum was thawed, and miR-122 was immediately measured in triplicate by PCR (median 1.7 pM (range 1.6-1.8)) and DCL (711 pM (686-737)). The samples were subsequently stored at room temperature and measured using both DCL assay and PCR. When the beads were added to the serum on day 0 the measurement of miR-122 by DCL remained constant until day 28, when there was an 18% loss. By contrast, when beads were not added, there was a substantial reduction in the measured concentration of miR-122 (day 28 miR-122 PCR: 1.3fM (0.7-1.8); DCL: 96pM (84-111)). The reduction in miR-122 was greater when PCR was used to measure miR-122, compared with bead addition/DCL on days 1-28 (Day 1 reduction in miR-122 concentration: PCR assay 58%, DCL assay 11%; day 7 reduction in miR-122 concentration: PCR assay 97%, DCL assay 43%).

**Figure 3.**
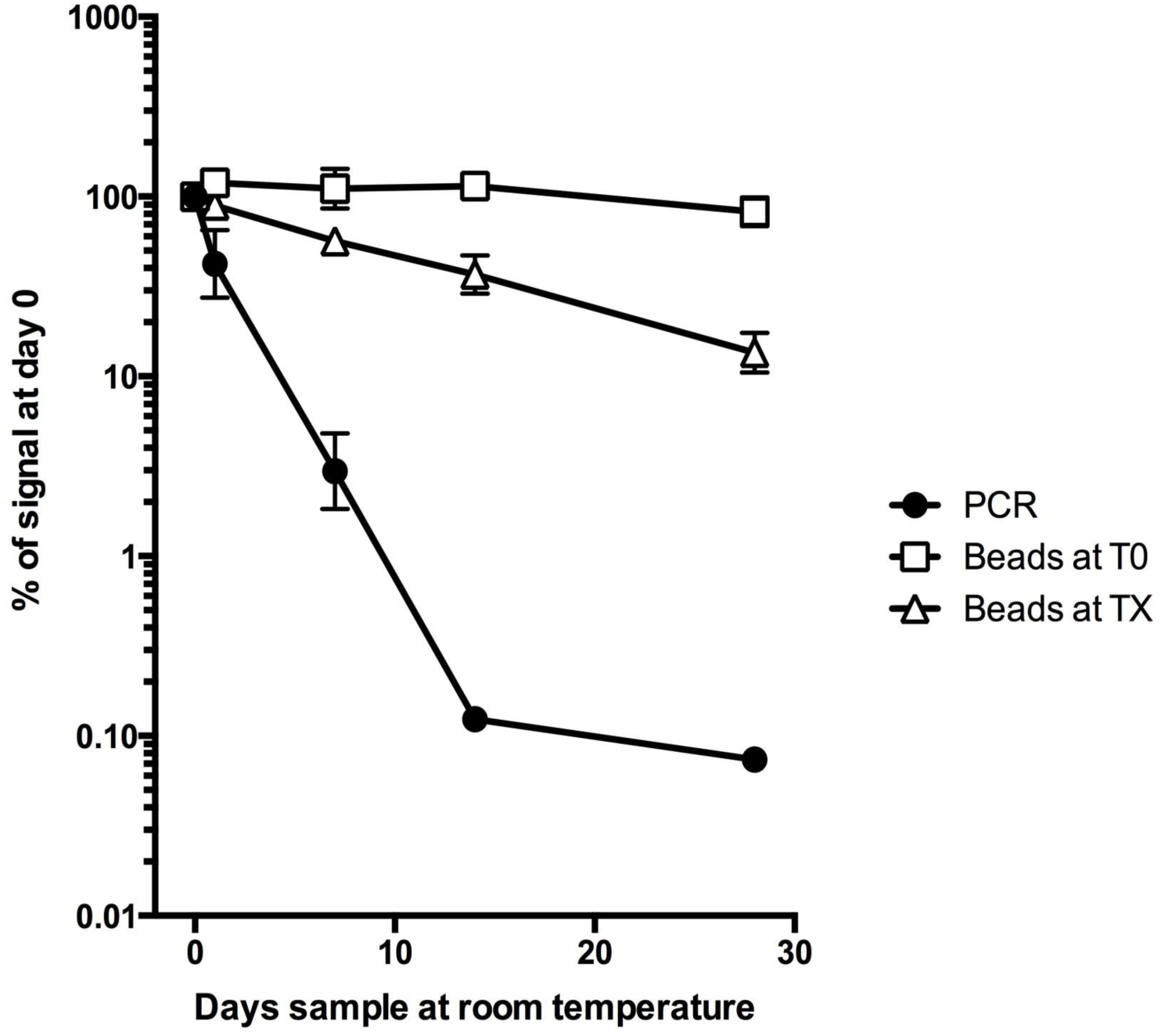
Degradation in the measurement of miR-122 is attenuated by adding DCL beads to serum. Serum from a patient with DILI had miR-122 measured by DCL assay and PCR at day 0. The serum was then stored at room temperature with beads added at day 0, and DCL assay was performed at days 1, 7, 14 and 28 (open square - beads at T0). Alternatively, the serum was stored at room temperature, and PCR (closed circle) or DCL (open triangle) was performed with the beads added on days 1, 7, 14 or 28 (beads at TX). The data are presented as the percentage of measurement at day 0 (mean, error bars represent 95%CI, N=3).

To understand the *ex vivo* kinetics of serum miR-122 degradation, we performed small RNA sequencing on healthy serum and serum from the patient with DILI (ALT > 1000 U/L after paracetamol overdose) at time 0 (immediately after thawing), 1 and 7 days after storage at room temperature. At time 0, miR-122 was present in high concentration in the form of multiple isomiRs in the DILI patient sample **(Figure 4).** The read counts are presented in **Supplementary Table 5.** The canonical form of miR-122 was not the most abundant in both health and DILI; miR-122_0_-1 was most common in the serum. This isomiR increased around 1100 fold with DILI to become the most abundant overall microRNA species in the circulation. Multiple other miR-122 isomiRs that had read counts less than 10 in the healthy serum increased substantially following DILI **(Supplementary Table 5).** The canonical miR-122 species decreased when sample processing was delayed with a concurrent increase in isomiRs, which is consistent with microRNA degradation. This degradation was faster in the serum from the patient with DILI compared to the healthy serum **(Supplementary Figure 4).**

**Figure 4.**
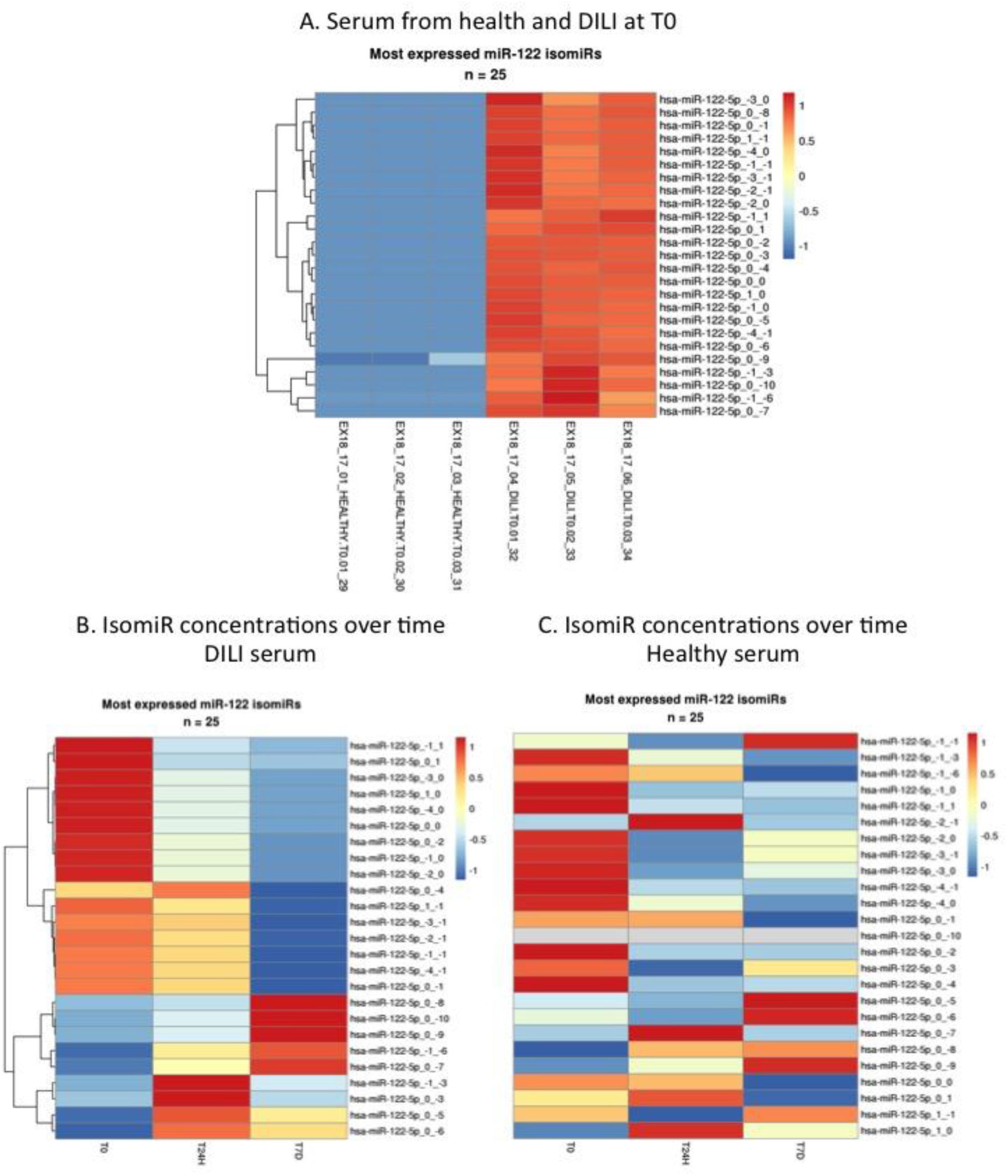
Heatmaps presenting the serum concentration of miR-122 isomiRs. A. Small RNA sequencing was performed on normal and DILI serum (ALT>1000U/L). Multiple isomiRs of miR-122 were increased with DILI when there was no delay in processing (T0) (N=3 per group). B. DILI serum was left at room temperature for 1 and 7 days (T24H and T7D). The change in isomiRs is presented. C. Healthy serum was left at room temperature for 1 and 7 days (T24H and T7D). The change in isomiRs is presented.

The presence of isomiRs may explain the discrepancy between miR-122 quantification by DCL and PCR assays. The effect of miR-122 isomiRs on quantification by PCR was determined using synthetic oligonucleotides corresponding to the canonical 22 base miR-122 human sequences and sequences with bases missing from the 3’ end (the most common isomiRs reported by sequencing in DILI). In both Taqman and SYBR-green based PCR efficiency was substantially reduced **(Figure 5).** The removal of a single base (the most common human miR-122 circulating species - hsa-miR-122-5p_0_-1) substantially reduced the efficiency of Taqman PCR (at 23 pM concentration Ct values: canonical miR-122: 19.5 (19.2 to 19.8), miR-122-5p_0_-1: 28 (28 to 28.2) P<0.0001 N=8). By contrast, with the DCL assay the efficiency was unaffected across all the isomiRs **(Figure 5).** To determine the impact of the presence of isomiRs on miR-122 analysis we created a ‘virtual patient’ using synthetic miR-122 isomiRs mixed in the same ratio as the RNA sequencing results to a final total microRNA concentration of 20 pM. Quantification by DCL produced a range of 21.7-28.1 pM across the isomiRs and the virtual patient. By contrast, with the virtual patient isomiR mix, Taqman-based PCR reported a concentration of 0.2 pM (0.19 to 0.21 N=6, using the miR-122_0_0 canonical standard curve) **(Figure 5).**

**Figure 5.**
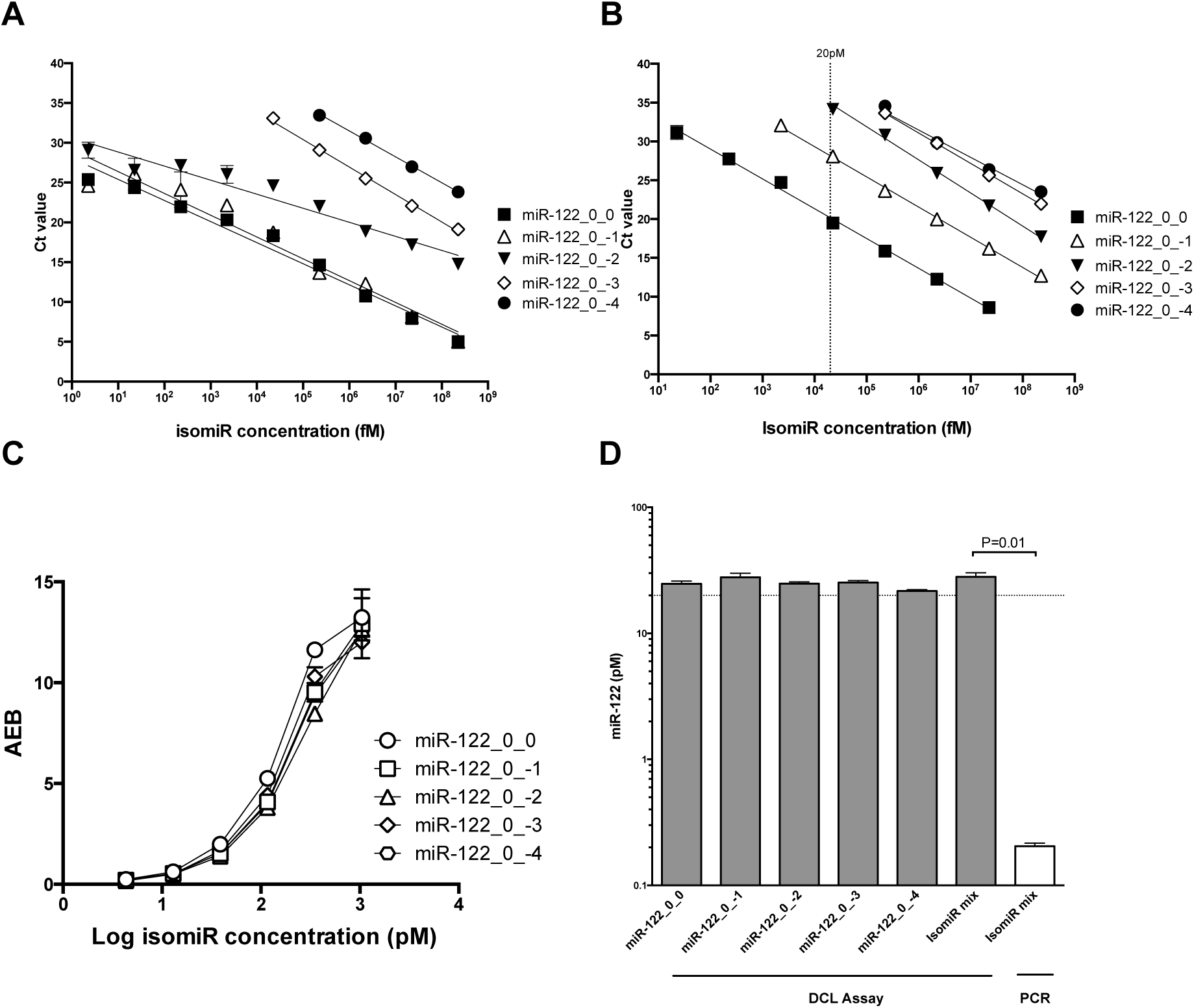
Standard curves for the quantification of miR-122 isomiRs by SYBR-green based PCR (A), Taqman based PCR (B) and DCL assay (C). AEB = Active Enzyme per Bead. D). A ‘virtual patient’ was created based on the sequencing results from Supplementary Table 5 using synthetic isomiRs (‘isomiR mix’). The final isomiR concentration for all groups was 20 pM (dotted line). The DCL assay was used to measure the concentration across individual isomiRs and the virtual patient isomiR mix. Taqman PCR was also used to quantify the microRNA concentration in the virtual patient using the miR-122_0_0 standard curve in panel B. In panel B 20 pM concentration is indicated by the dotted line. P value generated by Mann-Whitney Test (N = 6).

## Discussion

In this paper, we have described a new assay for the direct quantification of microRNA from serum and plasma without the need for microRNA extraction or amplification. Assay development revealed important insights regarding the biology of circulating microRNA that need to be carefully considered if researchers are to successfully qualify microRNAs as clinical biomarkers. We used miR-122 in the context of DILI as the clinical paradigm. Our data demonstrated that direct detection with the DCL assay identified DILI with high sensitivity and specificity that was comparable to the gold standard assay, PCR. In the presence of DILI, microRNAs were not stable *ex vivo* in serum and were present at high concentrations in the form of isomiRs. These isomiRs rendered analysis by commonly used ‘off-the-shelf’ PCR inaccurate. The DCL assay stabilised the microRNA target and prevented degradation, and accurately reported the total concentration of serum miR-122 isomiRs.

There has been a global research effort to develop microRNAs as disease biomarkers across a wide range of disease areas. However, to date, no microRNAs are used in routine clinical practice. The challenges to overcome include preventing degradation of target microRNA during sample processing and the development of rapid, user-independent, affordable and scalable assays suitable for use in standard clinical laboratories. The current gold standard assay is PCR, which is validated by decades of use and allows very sensitive quantification of microRNA. However, PCR requires skilled personnel, may be expensive for routine clinical use, and has multiple RNA extraction and amplification steps in which errors can occur. In our earlier publication we described a single probe method for detecting microRNA using single molecule arrays, with sequence specificity down to a single base, and without the use of amplification by polymerases.^7^ In the current paper, we built on this assay (the DCL assay) and demonstrate it can accurately measure microRNA directly in serum with minimal sample processing.

One of the most developed microRNA biomarkers is miR-122 in the context of DILI, especially secondary to paracetamol overdose. This clinical scenario is common – in the United Kingdom alone around 50,000 patients require emergency antidote treatment every year and, despite treatment, around 3 people die every week from paracetamol-induced acute liver failure.^17, 18^ Currently all patients deemed to need treatment receive the same dose of acetylcysteine, a practice which results in significant patient numbers being under or over treated.^19^ Once adopted into clinical practice, miR-122 promises to facilitate earlier decision making regarding both the stopping of unnecessary treatment and addition of new therapeutic agents currently being developed. In the current paper we analysed serum from 192 consecutive patients recruited to the MAPP Study and compared the quantification of miR-122 by DCL analysis to PCR. There was a significant correlation between miR-122 concentrations reported by PCR and DCL. Furthermore, subsequent DILI could be accurately predicted by DCL with high sensitivity and specificity that is comparable to the performance of PCR. These data support DCL being fit-for-purpose with regard to the stratification of patients after paracetamol overdose. An advantage of microRNAs as biomarkers is their sequence conservation across species. Rats and dogs are commonly used in pre-clinical drug development. miR-122 can be included as a biomarker of hepatocellular injury in short-term rat toxicology studies and its use adds benefit in detecting hepatotoxicity in this species.^20^ miR-122 is also a new biomarker for liver disease in dogs that is gaining traction in clinical veterinary practice.^16^ DCL reported miR-122 elevation in both these species. In summary, across species, DCL analysis accurately quantified miR-122 directly from serum or plasma without need for extraction or amplification. These properties meet our published target product profile for microRNA diagnostics in the liver injury space.^21^

The DCL assay uses an abasic PNA probe conjugated to superparamagnetic beads. We explored whether adding these beads to serum could prevent degradation of miR-122. We hypothesised that microRNA strands would hybridise to the abasic PNA probe forming a double strand PNA:RNA, hence being protected from both enzymatic^22^ and auto-hydrolysis degradation. The data demonstrate no loss of miR-122 signal over 14 days at room temperature. By contrast, there was a rapid decrease in signal when no beads were added (around 60% loss of PCR signal in the first day). This observation is directly contrary to the dogma that microRNAs are stable in the circulation but is consistent with other studies that have reported rapid degradation of miR-122 after sample collection.^23^ To explore the underlying biology we performed RNA sequencing on serum from DILI and healthy humans. This demonstrated that miR-122 isomiRs are present at high concentration in DILI – a finding that is consistent with previous studies. In healthy humans, miR-122 isomiRs have been demonstrated to be present in the circulation with a similar profile of types to our data. ^24^ Krauskopf *et al*. also used RNA sequencing and reported substantially increased circulating isomiRs with DILI in humans.^25^ In mice, Chowdhary *et al.* performed Northern blotting to demonstrate miR-122 isomiRs in the circulation following paracetamol toxicity.^26^ In our study, the concentration of isomiRs increased with sample degradation and, importantly, microRNA degradation was substantially increased with DILI.

The presence of circulating isomiRs for miR-122 has important consequences. Firstly, building on the work previously reported by Magee *et al.*,^27^ we demonstrated that both SYBR-Green and Taqman-based PCR is inaccurate due to variable efficiency across isomiRs. To demonstrate the clinical consequence we created a ‘virtual patient’ using synthetic isomiRs in the same ratio as reported by our sequencing of human DILI. The concentration of microRNA was 20 pM, with isomiRs mixed in the same ratio as DILI the reported PCR measured concentration was 0.2 pM, 100 fold less than the true value. The DCL assay reported the concentration of miR-122 canonical and isomiR sequences accurately (range 21.7-28.1pM). In the future development of miR-122 as a biomarker of liver toxicity, standard, ‘off the shelf’, PCR cannot be used a reliable assay. The relevance of isomiRs is not limited to DILI and miR-122 – for example, they are being actively developed as biomarkers for cancer. All researchers should consider whether their PCR assay is accurate in their experiments and the presence of isomiRs should be evaluated. Secondly, certain isomiRs are present at very high concentrations in DILI (for example, the isomiR lacking 3 bases from the 3’ end is the 9th highest abundance microRNA). The inaccuracy of PCR in the presence of variable isomiR concentrations may explain the inter-subject variability in miR-122 reported in the literature.^11^ The field should urgently re-evaluate the methods used in miR-122 quantification. Finally, microRNA degradation was more rapid in DILI than health. We speculate this effect may be because there is a release of RNA digesting enzymes from the liver into the circulation when injured and/or microRNA released from the injured liver is not protected from degradation by being bound to protein carriers or encapsulated in ECVs. A differential in degradation rate between health and disease has also been reported in patients with chronic kidney disease,^28^ and highlights the importance of robustly validated sample preparation protocols and approaches to prevent degradation.

There are limitations to the DCL assay. The limit of detection in 1.32 pM, which is higher than PCR (defined as 2.6 fM in our previous paper). ^7^ This limitation does not impact on the measurement of miR-122 because it is present in the circulation at relatively high abundance but may be a limiting factor for lower abundance microRNAs. The DCL assay as described in this manuscript measures all miR-122 isomiRs with equal efficiency. If there is an intellectual or commercial need then the assay probes could be designed to be selective for specific isomiRs.

In conclusion, successful translation of microRNAs from research tools to clinical assays needs a deeper understanding of their biology and assays that are fit-for-purpose. miR-122 is not as stable in serum *ex vivo* in DILI as health, and the presence of isomiRs make commonly used PCR methods unreliable. The developers of microRNA biomarkers should re-evaluate their PCR assays to determine if they are fit-for-purpose. The DCL assay can overcome these analytical and biological challenges and is a complementary tool to PCR.

