## Supplementary Data for "Direct detection of circulating microRNA-122 using dynamic chemical labelling with single molecule detection overcomes stability and isomiR challenges for biomarker qualification"

**Supplementary information**

**Stabilization study design**

<u>*Day 0 (T0)*</u>

**PCR analysis**:

a) Frozen serum samples were thawed and aliquoted for testing at T0 (day 0), T1 (day 1), T7 (day 7), T14 (day 14) and T28 (day 28). For each time point three replicates were used. (30 µL serum/replicate).

b) RNA extraction and PCR for T0 samples was performed using SYBR-green based PCR.

c) The remaining PCR samples were stored at room temperature.

**DCL analysis**:

a) Frozen serum samples were thawed and aliquoted for testing at T0, T1, T7, T14 and T28. For each time point three replicates were used. (30 µL serum/replicate).

b) For time point T0 three replicates were used (30 µL serum/replicate) containing serum + beads + Stabiltech buffer. T0 samples were analysed along with a calibration curve.

c) For time points T1, T7, T14, T28 there were two groups of samples:

Group 1. Serum + beads + Stabiltech Buffer at T0. Analysis was performed at performed at T1, T7, T14 or T28.

Group 2. Serum kept at room temperature until addition of beads + Stabiltech buffer at T1, T7, T14 or T28. Analysis was performed at T1, T7, T14 or T28.

<u>*Day 1 (T1), Day 7 (T7) , Day 14 (T14) and Day 28 (T28).*</u>

**PCR analysis**:

RNA extraction and SYBR-green based PCR was performed for room temperature stored T1, T7, T14 or T28 samples.

**DCL analysis**:

a) Beads + lysis buffer were added to the samples from Group 2 (T1, T7, T14 or T28). Calibration curves were prepared at the same time as the combining of beads plus lysis buffer with the samples.

b) Samples from Groups 1 and 2 (T1, T7, T14 or T28) were analysed along with the corresponding calibration curves.

**Supplementary Table 1.** Patient demographics from MAPP Study.
**Supplementary Table 2.** Data from rat model of paracetamol toxicity.
**Supplementary Table 3.** Calibrator concentrations calculated by DCL assay and ddPCR.
**Supplementary Table 4.** Clinical details of dogs with liver disease.
**Supplementary Table 5.** miR-122 isomiRs in health and DILI serum. RPM = reads per million.
**Supplementary Table 6.** Sequences of capture probe **(1)** and miR-122 target **(2)**.

| | |
|---|---|
| Number | 192 |
| Sex (number M:F) | 66:126 |
| Age (years) | 28 (19-39) |
| Amount of paracetamol ingested (g) | 17 (9-21) |
| Time from ingestion to first blood sample (h) | 4 (4-8) |
| Admission paracetamol concentration (mg/L) | 111 (62-150) |
| Admission ALT (U/l) | 16(12-24) |
| Admission ALP (U/l) | 69 (46-78) |
| Admission INR | 1 (1-1.1) |
| Admission serum creatinine (μmol/L) | 59 (50-68) |
| Number with admission ALT <ULN | 175 |
| Number with admission ALT >100 | 3 |
| Number with admission ALT >1000 | 0 |
| Number with Peak ALT >100 | 18 |
| Number with Peak ALT >1000 | 9 |

**Supplementary Table 1.** Patient demographics from the MAPP Study. ALT = serum alanine transaminase activity. ALP= serum alkaline phosphatase activity. INR= international normalised ratio**.** Data are presented as median (IQR) for continuous variables.

| Animal N° | Group | Dose (mg/kg) | Hepatocyte Necrosis Score | Hepatocyte Apoptosis Score | ALT (U/L) | GLDH (U/L) | DCL miR-122 (pM) | DCL miR-122 (copy/µL) | ddPCR miR-122 (copy/µL) |
|---|---|---|---|---|---|---|---|---|---|
| 61 | Control | 0 | 0 | 0 | 30 | < LLOQ | < LLOQ | < LLOQ | 1.2E+03 |
| 62 | Control | 0 | 0 | 0 | 23 | < LLOQ | 14 | 8.4E+06 | 1.2E+03 |
| 63 | Control | 0 | 0 | 0 | 91 | < LLOQ | < LLOQ | < LLOQ | 4.5E+02 |
| 64 | Control | 0 | 0 | 0 | 21 | < LLOQ | < LLOQ | < LLOQ | 1.5E+03 |
| 65 | Control | 0 | 0 | 0 | 20 | 6 | < LLOQ | < LLOQ | 3.3E+03 |
| 66 | Control | 0 | 0 | 0 | 30 | 6.4 | 18 | 1.1E+07 | 5.6E+03 |
| 67 | Paracetamol | 1500 | 1 | 4 | 674 | 119.3 | 2705 | 1.6E+09 | 2.0E+05 |
| 68 | Paracetamol | 1500 | 0 | 2 | 76 | 62.7 | 435 | 2.6E+08 | 5.7E+04 |
| 69 | Paracetamol | 1500 | 0 | 1 | 41 | 5.5 | 32 | 1.9E+07 | 1.1E+04 |
| 70 | Paracetamol | 1500 | 4 | 3 | 1985 | 431.9 | 5345 | 3.2E+09 | 1.4E+06 |
| 71 | Paracetamol | 1500 | 0 | 2 | 184 | 234.1 | 594 | 3.6E+08 | 2.8E+05 |
| 72 | Paracetamol | 1500 | 0 | 1 | 153 | 392.4 | 1056 | 6.4E+08 | 4.5E+05 |

**Supplementary Table 2.** Data from rats treated with a single dose of paracetamol (1500mg/kg oral) or vehicle control. Plasma miR-122 concentrations were measured by both DCL and ddPCR. Hepatocellular necrosis and apoptosis were scored on H&E-stained sections of the right lateral liver lobe as described in the methods. Hepatocellular apoptosis was further confirmed by immunohistochemical detection of activated caspase 3 on formalin fixed, paraffin-embedded sections of the liver. LLOQ – lower limit of quantification.

| Theoretical titer | | | | |
|---|---|---|---|---|
| **Study ID** | **Copy/µL** | **pM** | | |
| C-1 | 6022000 | 10 | | |
| C-2 | 60220000 | 100 | | |
| C-3 | 301100000 | 500 | | |
| **ddPCR evaluation** | | | | |
| **Study ID** | **Copy/µL** | **pM** | **Fold difference expected vs obtained** | **CV (%)** |
| C-1 | 2918 | 0.0048 | 2083.3 | 4.2 |
| C-2 | 11517 | 0.0191 | 5235.6 | 7.2 |
| C-3 | 28746 | 0.0477 | 10482.1 | 3.9 |
| **DCL evaluation** | | | | |
| **Study ID** | **Copy/µL** | **pM** | **Fold difference expected vs obtained** | **CV (%)** |
| C-1 | 4145513 | 6.883 | 1.45 | 16.8 |
| C-2 | 43032534 | 71.43 | 1.39 | 2.2 |
| C-3 | 212409632 | 352.66 | 1.41 | 8.0 |

**Supplementary Table 3.** Calibrator concentrations of miR-122 spiked into healthy rat plasma obtained by both the DCL assay and ddPCR. Fold differences between expected calibrator concentrations and those obtained are expressed as the ratio between expected vs obtained values. CV = coefficient of variation.

| Breed | Age (Yrs) | Diagnosis | ALT (U/L) |
|---|---|---|---|
| Havanese | 8 | cholestasis | 656 |
| English Springer Spaniel | 11 | chronic hepatitis | 193 |
| Scottish Terrier | 9 | lymphoma | 233 |
| Flat Coated Retriever | 9 | adenoma | 776 |
| English Pointer | 6 | granuloma, hepatitis | 1544 |
| Cairn Terrier | 11 | hepatocellular carcinoma | 182 |
| Labrador | 8 | chronic-active hepatitis | 295 |
| West Highland White Terrier | 2 | none | 55 |
| Old English Sheepdog | 5 | hepatitis, chronic | 1235 |
| Labrador Cross | 13 | cholangiocellular carcinoma | 46 |
| Bearded Collie | 13 | hepatopathy | 33 |
| Labrador | 8 months | lobular dissecting hepatitis | 201 |

**Supplementary Table 4.** Clinical details of dogs with liver disease.

| IsomiR | Healthy sample 1 (RPM) | Healthy sample 2 (RPM) | Healthy sample 3 (RPM) | DILI sample 1 (RPM) | DILI sample 2 (RPM) | DILI sample 3 (RPM) |
|---|---|---|---|---|---|---|
| hsa-miR-122-5p_0_-1 | 187.85 | 366.56 | 274.7 | 332446.44 | 299487.41 | 311780.78 |
| hsa-miR-122-5p_0_0 | 95.89 | 151.16 | 137.72 | 142689.37 | 138157.28 | 137990.22 |
| hsa-miR-122-5p_0_1 | 89.56 | 68.4 | 81.62 | 42105.79 | 44347.71 | 44885.22 |
| hsa-miR-122-5p_0_-3 | 2.66 | 14.22 | 6.83 | 9849.26 | 9791.23 | 9543.85 |
| hsa-miR-122-5p_0_-2 | 1.9 | 14.62 | 4.01 | 8384.43 | 8419.18 | 8315.69 |
| hsa-miR-122-5p_-1_-1 | 1.27 | 3.89 | 3.86 | 6194.05 | 5296.08 | 5674.95 |
| hsa-miR-122-5p_-1_0 | 1.14 | 1.21 | 1.04 | 2698.76 | 2540.07 | 2458.8 |
| hsa-miR-122-5p_0_-5 | 1.39 | 1.21 | 1.63 | 2173.99 | 2008.81 | 1959.81 |
| hsa-miR-122-5p_0_-6 | 0.25 | 7.24 | 9.2 | 2171.1 | 2149.94 | 2041.44 |
| hsa-miR-122-5p_0_-4 | 1.39 | 0.94 | 1.48 | 1977.07 | 1897.31 | 1951.41 |
| hsa-miR-122-5p_-2_-1 | 0.38 | 0.67 | 1.34 | 1367.41 | 1166.56 | 1192.23 |
| hsa-miR-122-5p_-4_-1 | 1.01 | 0.54 | 1.19 | 1109.43 | 1063.8 | 1002.78 |
| hsa-miR-122-5p_-1_1 | 0.38 | 0.13 | 0.15 | 1016.45 | 1072.23 | 1148.35 |
| hsa-miR-122-5p_1_-1 | 0.25 | 0.4 | 0.45 | 910.26 | 830.8 | 849.17 |
| hsa-miR-122-5p_-3_-1 | 0.25 | 0.54 | 0.15 | 777.5 | 658.08 | 698.09 |
| hsa-miR-122-5p_-2_0 | 0.13 | 0.13 | 0.15 | 455.45 | 402.66 | 400.06 |
| hsa-miR-122-5p_-4_0 | 0.13 | 0.13 | 0.15 | 419 | 345.62 | 375.48 |
| hsa-miR-122-5p_1_0 | 0 | 0.27 | 0.15 | 359.01 | 337.09 | 333.9 |
| hsa-miR-122-5p_-3_0 | 0.13 | 0 | 0.3 | 269.64 | 210.18 | 238.94 |
| hsa-miR-122-5p_-1_-3 | 0 | 0.13 | 0.3 | 176.78 | 206.82 | 170.2 |
| hsa-miR-122-5p_0_-7 | 0.13 | 0 | 0 | 116.41 | 121.28 | 100.75 |
| hsa-miR-122-5p_0_-8 | 0 | 0.27 | 0 | 89.5 | 81.15 | 85.5 |
| hsa-miR-122-5p_0_-9 | 0 | 0 | 16.62 | 72.08 | 79.82 | 77.27 |
| hsa-miR-122-5p_-1_-6 | 0 | 0 | 0.15 | 46.24 | 53.86 | 39.21 |
| hsa-miR-122-5p_0_-10 | 0 | 0 | 0 | 29.42 | 35.9 | 30.68 |

**Supplementary Table 5.** miR-122 isomiRs in health and DILI serum. RPM = reads per million.

| ID | Name | Peptide with abasic position (N'-C') |
|---|---|---|
| **1** | Capture probe | xx-CACCATT*GT *GL* AC*ACT*CCA |
| | | **miRNA-122-5p sequence (5'-3')** |
| **2** | Target miR-122 | *UGGAGUGU**G**ACAAUGGUG*UUUG |

**Supplementary Table 6.** Sequences of capture probe **(1)** and miR-122 target **(2)**.
Key: xx = amino-PEG linker; T* = thymidine containing a propanoic acid side chain at the gamma position; C* = cytosine containing a propanoic acid side chain at the gamma position; *GL* = abasic "blank" monomer presenting a secondary amine and containing a propanoic acid side chain at the gamma position; the italicized bases in **2** form a duplex with the capture probe, and G at position 9 (bold) is opposite the "blank" monomer and binds to the aldehyde-modified cytosine. The mature sequence of miRNA-122 is 22 bases long. Our probe targets 18 bases out of the 22, leaving out the 4 bases at the 3'-end.

**Supplementary Figure 1.** miR-122 measured by DCL and PCR in serum samples from the MAPP study.
**Supplementary Figure 2.** Correlation between miR-122 concentrations of rat plasma samples (see Supplementary Table 2) calculated by ddPCR and DCL.
**Supplementary Figure 3.** DCL assay quantifies increased circulating miR-122 in dogs
**Supplementary Figure 4.** Degradation of miR-122 isomiRs over time.
**Supplementary Figure 5.** Dynamic chemistry labeling of biotin into an immobilized abasic PNA probe on a superparamagnetic bead.

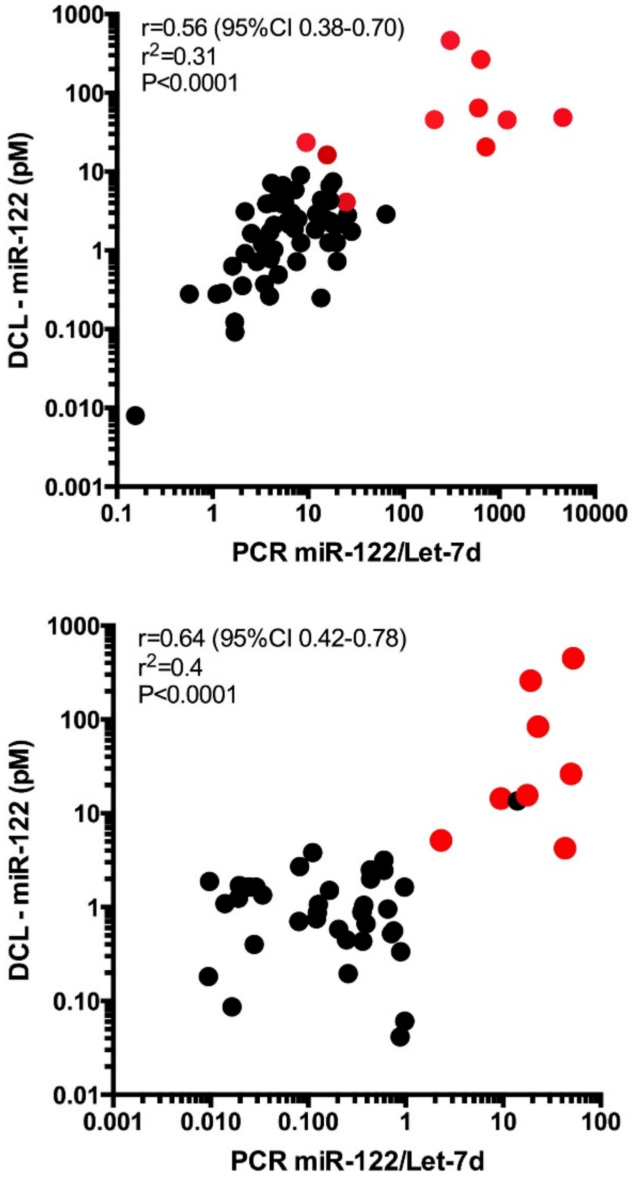


**Supplementary Figure 1.** miR-122 was measured by DCL and PCR in serum samples from the MAPP study. The PCR data is from the original analysis with normalisation to the microRNA Let-7d. The two panels present data from patients recruited at two UK hospitals, St Thomas' Hospital London and Aberdeen Royal Infirmary. Pearson correlation was performed to explore the relationship between the two techniques. Red points represent study participants who developed acute liver injury (ALT > 100U/L).

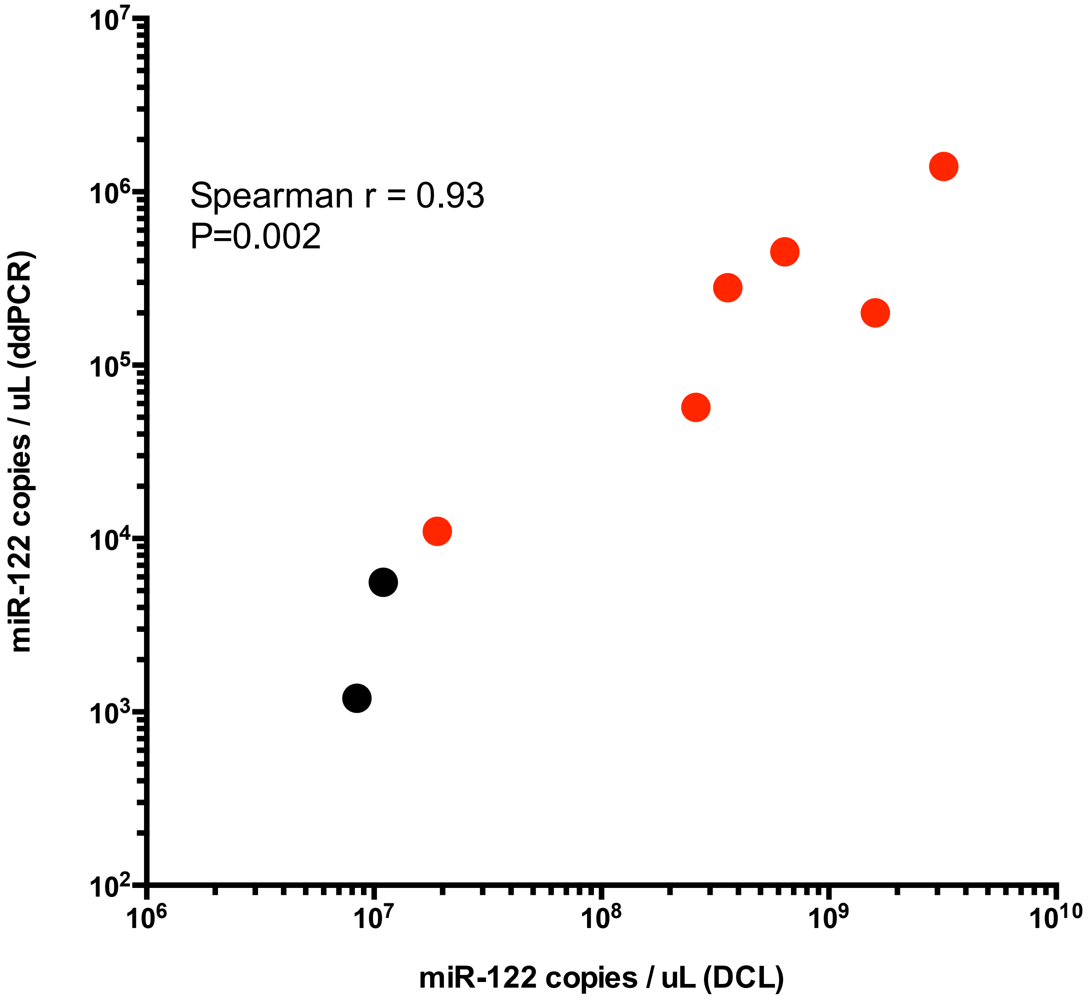


**Supplementary Figure 2.** Correlation between miR-122 concentrations of rat plasma samples (see Supplementary Table 2) calculated by ddPCR and DCL. Red dots are rats treated with paracetamol. The two black dots correspond to the 2 control rats with values of DCL above LLOQ. The values of ddPCR in the 4 control rats with DCL values below the LLOQ (not represented in the graph) were between 4.5 E+02 and 3.3 E+03 copies per microliter.

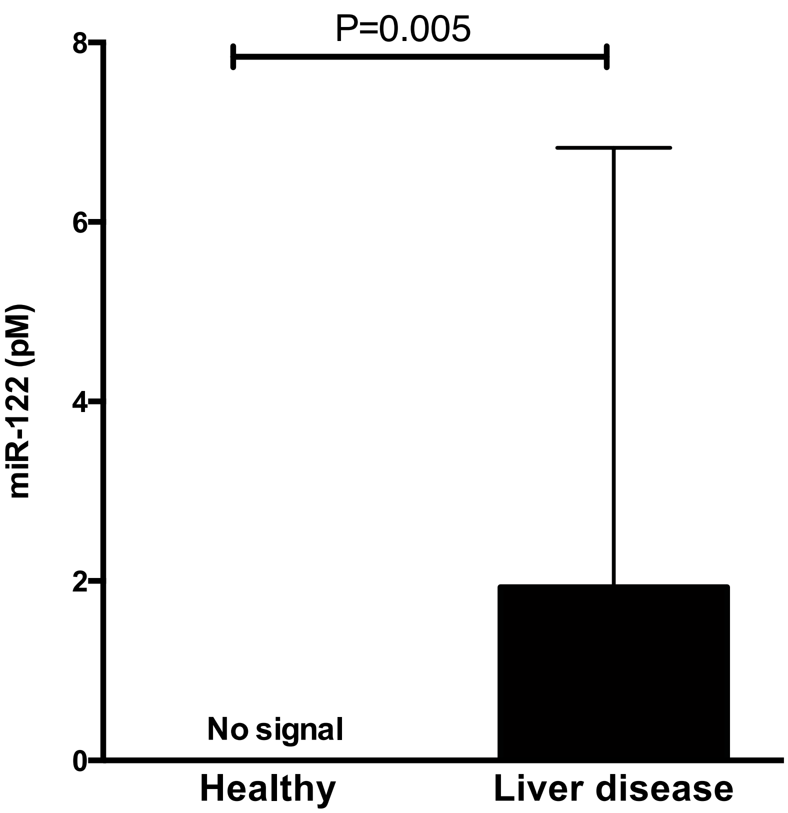


**Supplementary Figure 3.** DCL assay quantifies increased circulating miR-122 in dogs. miR-122 was increased in dogs with clinical liver disease compared with healthy dogs (N=12 per group). Graphs present median values, bars present the IQR. P values calculated by Mann-Whitney test.

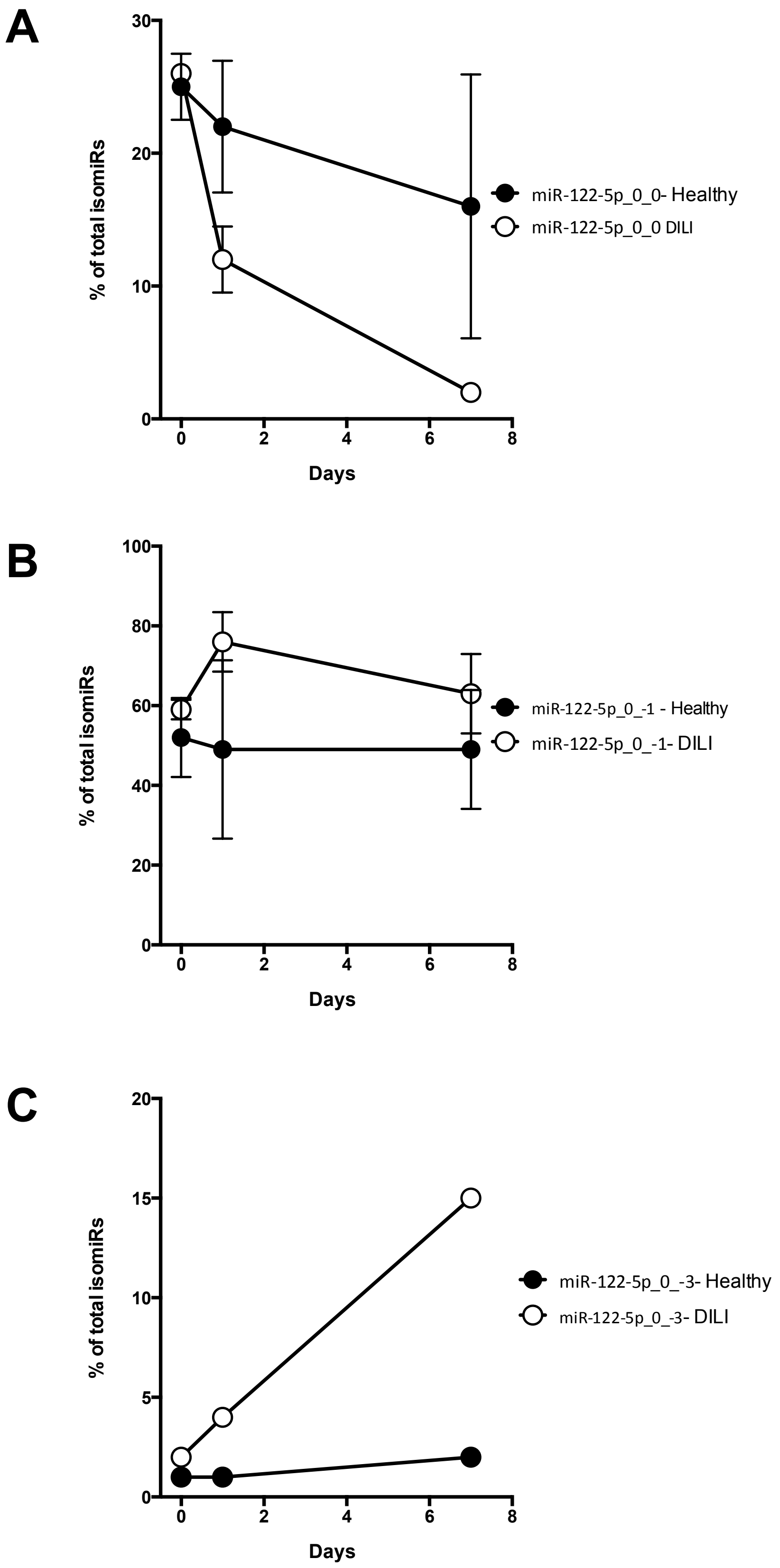


**Supplementary Figure 4.** RNA sequencing demonstrated that the canonical form of miR-122 (miR-122-5p_0_0) (A) degrades over time with a concurrent increase in shorter isomiRs (miR-122-5p_0_-1 and miR-122-5p_0_-3) (B and C, repectively). The degradation is enhanced with DILI. Data are represented as mean and 95%CI.

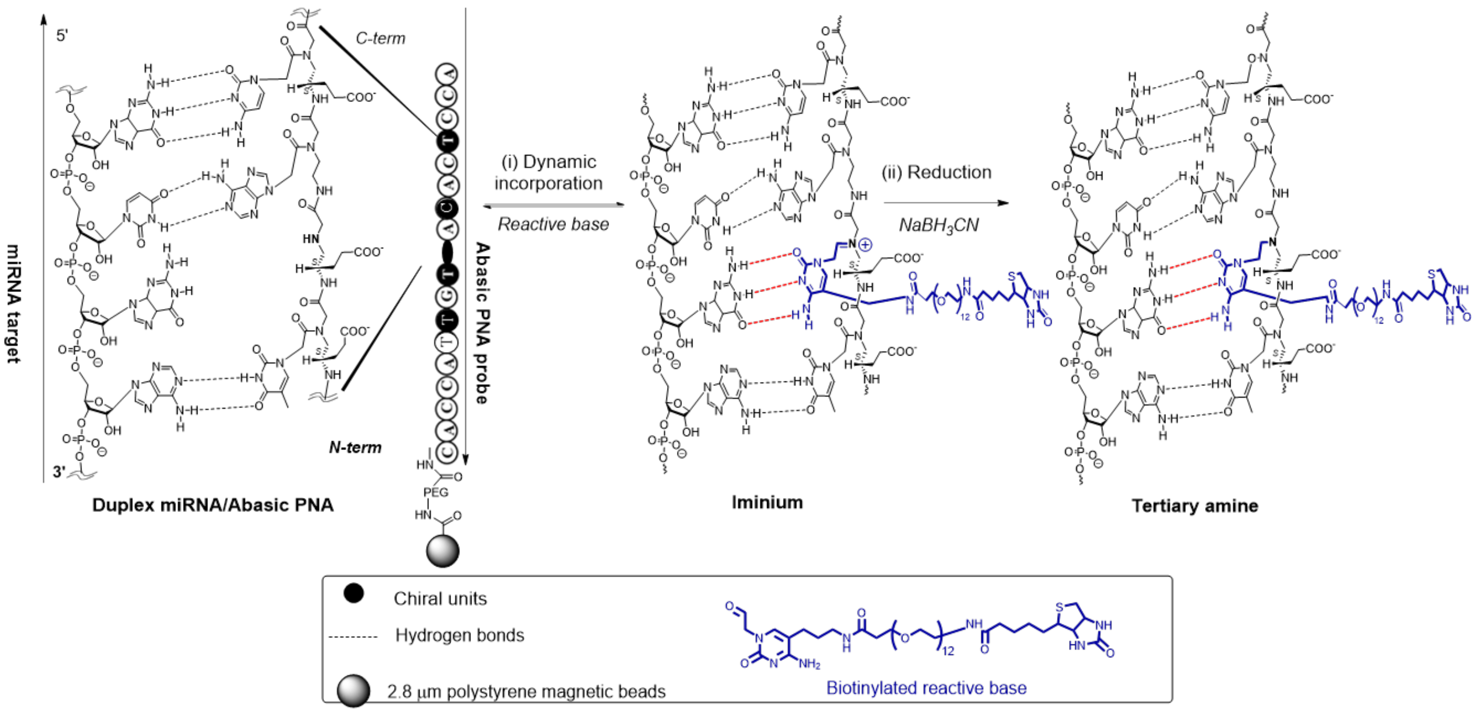


**Supplementary Figure 5.** Dynamic chemistry labeling of biotin into an immobilized abasic PNA probe on a superparamagnetic bead.
